# BEEP Learning: Multi-View Image Decomposition for Massively Multiplexed Biological Fluorescence Microscopy

**DOI:** 10.64898/2026.02.19.706833

**Authors:** Ruogu Wang, Thet Teresa Hnin, Yunlong Feng, Alex M. Valm

## Abstract

Fluorescence imaging with spectrally variant fluorophores allows the spatial mapping of biological structures with exquisite cellular and molecular specificity. However, the ability to robustly discriminate multiple fluorophores in any single imaging experiment is greatly hindered by the broad emission spectra of bio-compatible fluorophores and the large contribution of noise in low-energy regime fluorescence microscopy. In this study, we propose a novel machine learning framework, Bleaching-Excitation-Emission Photodynamics (BEEP) learning, that exploits multiple discriminatory features of fluorescent dyes to greatly expand the number of distinguishable objects in an image by integrating emission spectra, excitation variability, and bleaching dynamics into a unified multi-view, fluorescence unmixing approach. Our method is built upon a rank-one-tensor-based generalized linear model and leverages two biophysically grounded assumptions: consistent spectral and bleaching behaviors under fixed excitation, and invariant fluorophore abundances across excitations. We first extract excitation-specific spectral and bleaching signatures from reference images, and then use them to estimate abundances in complex mixtures. Experimental results on both simulated and real images of microbial populations demonstrate that our approach significantly outperforms conventional and partially multi-view methods, offering improved robustness and accuracy in highly multiplexed fluorescence imaging.

## 1 Main

Modern biological research increasingly demands a holistic understanding of cellular environments, where the spatial coordination of diverse molecular species underlies function Hartwell et al (1999); Kitano (2002); Moffitt et al (2022); Moses and Pachter (2022). Multiplex fluorescence microscopy has emerged as a transformative tool across biological disciplines, that eschews observation of isolated biomolecules for visualization of complex cellular and molecular interactomes within and among cells Gerdes et al (2013); Valm et al (2016); Tran et al (2025). While tens to hundreds of spectrally variant fluorescent reporter molecules have been developed and employed individually or in low complexity bioimaging scenarios, the simultaneous use of multiple fluorophores is severely hampered by their broad and overlapping emission spectra Wysocki and Lavis (2011); Waters (2009).

To circumvent the problem of spectral overlap, fluorescence microscopy detectors may acquire many images over discretely sampled wavelength bands Garini and Tauber (2012); Li et al (2013). State-of-the-art approaches extract the spectral image patterns of known fluorophores from labeled datasets then utilize least squares-based regression to assign the class and abundance of these fluorophores to real biological images acquired under the same conditions as the known labeled data; a framework known in the bioimaging field as linear unmixing Zimmermann (2005a); Neher et al (2009). However, the inherent uncertainty in sampling photons in the low energy regime of molecular fluorescence, i.e., shot noise, combined with photobleaching, the unavoidable loss of signal that occurs over time while fluorophores suffer non-reversible molecular rearrangements in the excited state, set practical limits on the narrowness and number of wavelength bands that can be acquired in any single imaging experiment, and therefore the separability of different fluorescent signals when applying linear unmixing to multispectral fluorescence images Waters (2009); Kalies et al (2011).

To address this shortcoming and therefore increase the number of different fluorophores that can be distinguished with suitable confidence in biological fluorescence images, we introduce a flexible linear unmixing model that comprehensively accounts for the variability introduced by overlapping emission and excitation spectra when images are acquired over diverse excitation conditions and in the context of dynamic photobleaching. We introduce a novel rank-one-tensor-based generalized linear unmixing model and its application to decompose images acquired with specialized methods that we call Bleaching-Excitation-Emission Photodynamics (BEEP) learning, that for the first time, simultaneously incorporates fluorescence bleaching, excitation, and emission photodynamics within a unified unmixing process. Because spectrally variant fluorophores have characteristic excitation spectra and unique bleach profiles, in addition to equally characteristic emission spectra that can be learned from microscope images, our approach turns the challenges described above into opportunities, leading to enhanced signal separation and more accurate fluorophore quantification.

The concept of leveraging complementary information from multiple data views is well-established in machine learning and has led to the development of multiview approaches that demonstrate improved robustness, flexibility, and generalization in various applications Hotelling (1992); Muslea et al (2000); Sun (2013); Xu et al (2013); Zhao et al (2017). In the context of biological spectral unmixing, image acquisition and fluorophore estimation methods that leverage excitation and emission have shown promise in enhancing the accuracy of fluorophore discrimination and abundance estimation Valm et al (2016); Neher et al (2009); Lakowicz (2006); Chen et al (2022); Gautheron et al (2023). Building on these successes, we previously developed a multi-view learning framework that incorporates discreetly sampled emission spectra obtained with varying excitation wavelengths to enhance the quantification of fluorophores with highly overlapping spectra and increase the number of distinguishable fluorescent labels for microbiome imaging Wang et al (2025b).

Separately, photobleaching kinetics have been explored for distinguishing fluorophores in biological imaing. Photobleaching is a phenomenon characterized by a gradual reduction in emission intensity from a fluorescent sample due to prolonged exposure to light, which leads to irreversible chemical modifications within the fluorophore molecule Demchenko (2020). The emission intensity of an ensemble of fluorophore molecules follows an exponential decay pattern Pawley (2006). Notably, fluorophores with identical spectral characteristics can exhibit distinct and highly characteristic bleaching rates due to differences in their chemical and physical environments Song et al (1995). This bleaching-rate variability, when combined with spectral differences, offers a powerful basis for distinguishing and quantifying fluorophores with highly overlapping emission spectra, thereby improving spectral unmixing and abundance estimation across multiple biological applications Brakenhoff et al (1994); Orth et al (2018); Hugelier et al (2021); Wüstner (2022).

In this study, we propose a novel multi-view machine learning framework, BEEP, for Bleaching-Excitation-Emission-Photodynamics Learning that integrates bleaching curves as well as excitation and emission spectra for robust fluorophore unmixing, Figure 1. Our method begins by acquiring image sequences through repeated imaging of the sample under various excitation wavelengths to intentionally induce photobleaching. This design enables us to capture the dynamic response of fluorophores under diverse excitation conditions, yielding a rich, multi-dimensional dataset for unmixing.

**Fig. 1.**
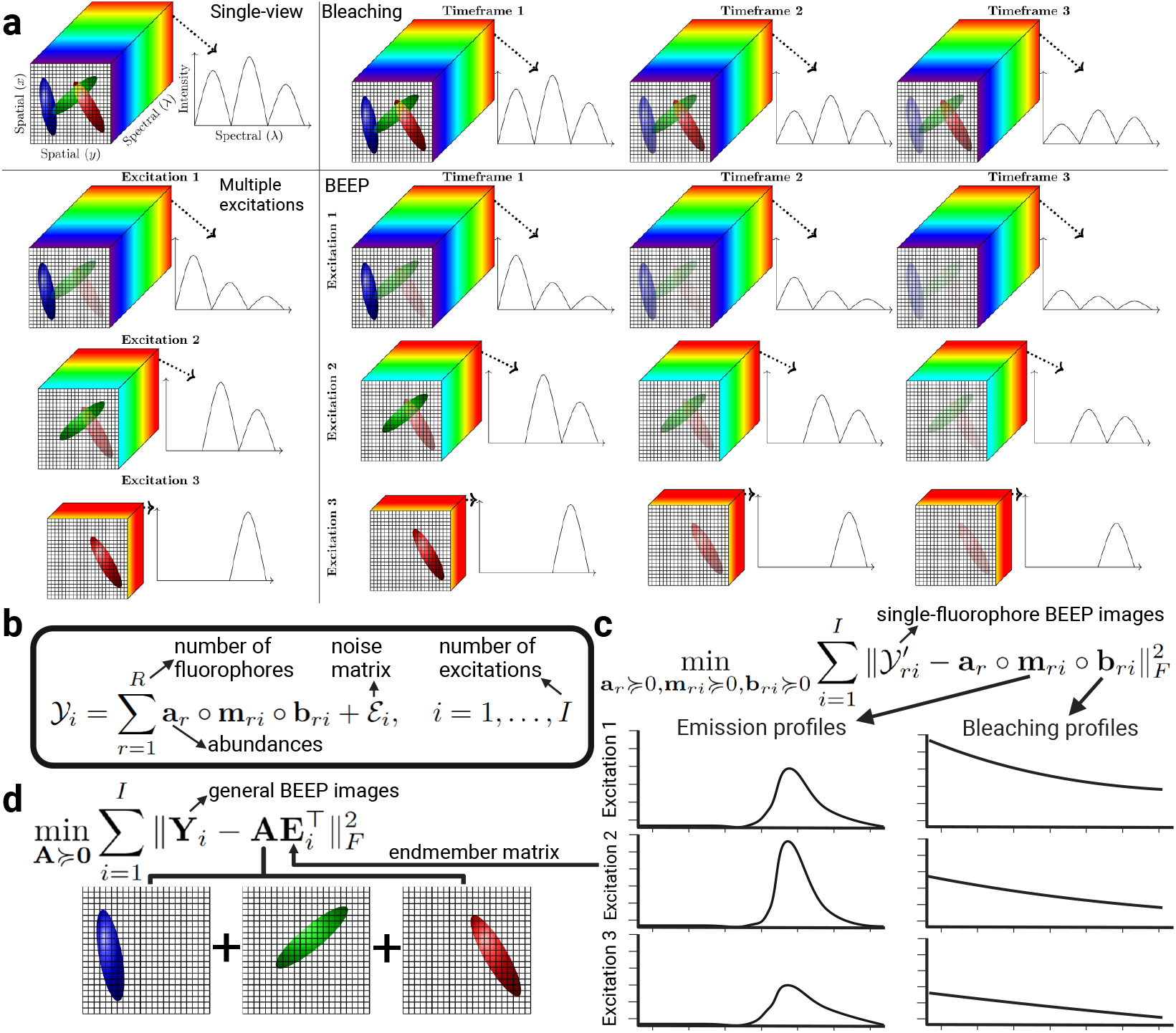
Overview of the biological spectral unmixing workflow and BEEP learning process. (a) Different data modalities used in the analysis: single-view data (emission spectra), photobleaching dynamics data (temporal intensity decay), multiple excitation data (emission spectra under varying excitation wavelengths), and BEEP data (integrated bleaching, excitation, and emission photo dynamics). (b) Tensor-based linear mixture model. (c) BEEP learning for endmember extraction. (d) BEEP learning for abundance estimation.

The foundation of our approach rests on two key assumptions. First, while a fluorophore’s emission spectral signature and bleaching rate may vary across different excitation conditions, they remain consistent within each specific excitation condition. This within-condition consistency allows us to reliably model each fluorophore’s behavior and use it to disentangle mixed signals. Second, the abundance of each fluorophore molecule remains constant across excitation conditions, as excitation settings during imaging do not alter the sample’s molecular concentration. These biologically reasonable assumptions enable us to leverage the dynamic, multi-view information captured during imaging of fixed biological samples to perform accurate fluorophore quantification.

Our BEEP learning framework proceeds in two stages. First, it learns excitation-specific spectral and bleaching signatures from reference image sequences. Next, it uses these learned signatures to estimate fluorophore abundances in new samples imaged under the same experimental conditions. By incorporating spectral and temporal bleaching information, our method resolves overlapping signals more effectively than traditional or limited-view methods.

To evaluate the performance of our approach, we conducted experiments on both simulated datasets and real microbial mixture images. Our results show that BEEP outperforms conventional single-view methods, as well as two-view methods that combine excitation and emission spectra or bleaching and emission spectra. These results underscore the benefits of incorporating bleaching dynamics into the unmixing framework, which provides an additional orthogonal dimension of information that significantly improves the accuracy and robustness of fluorophore separation and quantification.

## 2 Results

In the following experiments, we compare the performance of BEEP learning with three approaches: (1) single-view spectral unmixing Zimmermann (2005b), (2) multiview learning with photobleaching and emission spectra Hugelier et al (2021), and (3) multi-view learning with excitation and emission spectra Wang et al (2025b). Figure 1b illustrates the single-view and multi-view data for each method.

### 2.1 Simulations

In this section, we utilize simulated data to demonstrate the effectiveness of our proposed method in addressing the challenges of spectral overlap in hyperspectral imaging. To achieve this, we generate a comprehensive dataset that incorporates emission spectra, excitation spectra, photobleaching dynamics, and realistic noise conditions, enabling a robust evaluation of our method’s performance.

To assess the capability of our approach in handling highly overlapping endmembers across varying numbers of views, we generated 100 skewed Gaussian-distributed emission spectra and excitation spectra, each with 32 channels, to recapitulate the skewed spectra of widely used organic fluorophores imaged with commercially avaialble spectral confocal microscopes Fig 2(a) and (b). The correlation coefficient between any two adjacent endmembers is at least 0.97, reflecting a high degree of spectral overlap and simulating the challenging conditions that would be encountered in real-world scenarios when 100 organic fluorophores with emission spectra constrained within the visible to near-infrared wavelength range are used for light microscopy.

**Fig. 2.**
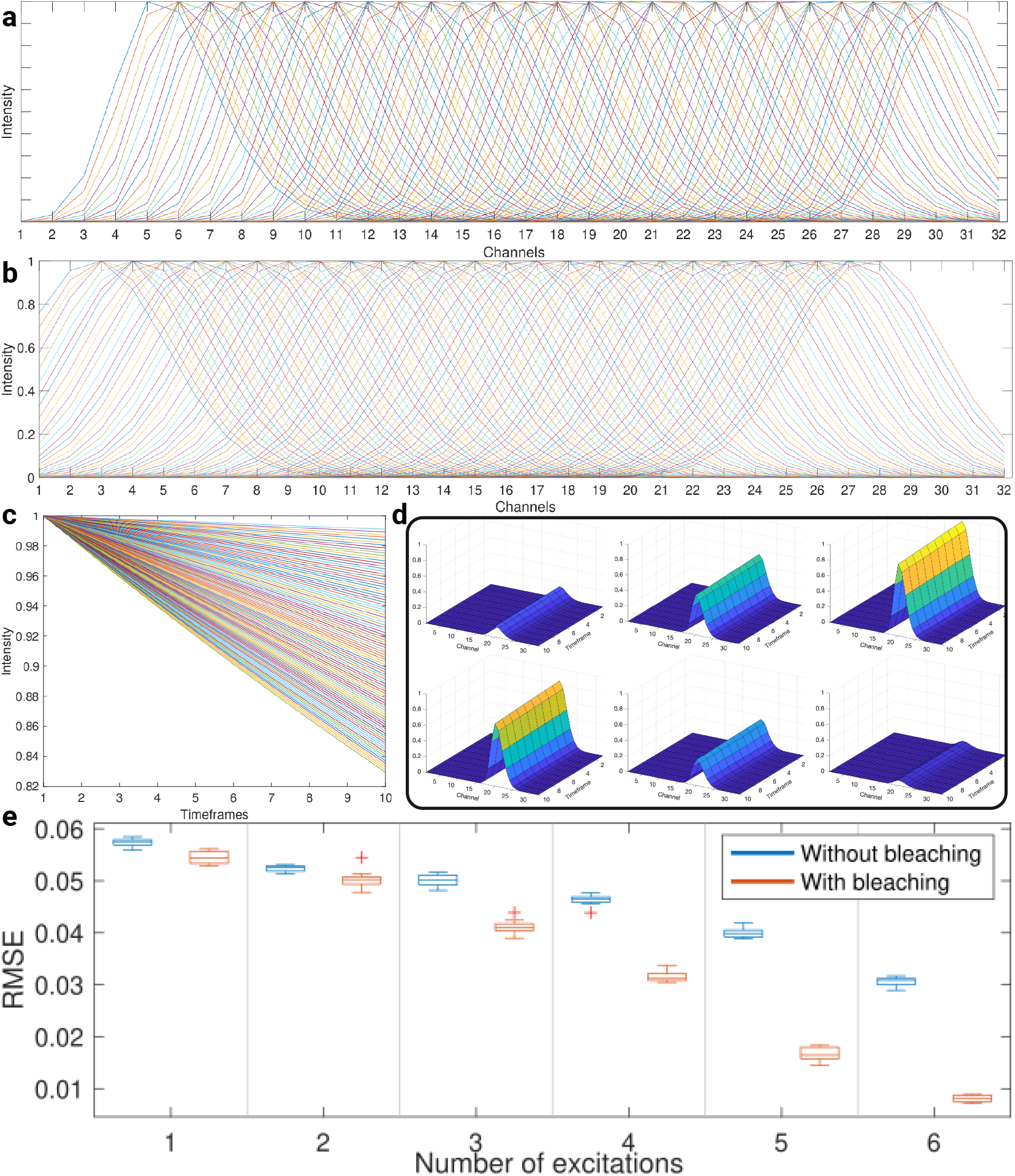
Simulation results. (a) 100 simulated emission spectra. (b) 100 simulated excitation spectra. (c) 100 bleaching profiles. (d) Example simulated BEEP signature. (b) Boxplots of Root Mean Square Errors (RMSEs) comparing abundance estimation accuracy for simulations with varying numbers of excitations. Each RMSE represents the error between the estimated and true abundances in simulated spectral images.

Next, we generate 100 photobleaching profiles following a negative exponential distribution, with bleaching rates ranging from 0.001 to 0.02, Figure 2(c).This range captures the observed variability in photobleaching dynamics in real-world settings, ensuring that our simulations reflect the diverse behaviors of fluorophores under different conditions in a biolophysically meaningful way.

Using these profiles, we derive the BEEP signature, as illustrated in Figure 2(d). Subsequently, we simulate 1000 spectral images, each comprising 1000 pixels. For each pixel, we randomly assign the abundance of one fluorophore from a uniform distribution *U* (0, 1), while setting the abundances of the remaining fluorophores to zero. This setup mimics the sparsity often observed in real-world samples, where only a subset of fluorophores may be present in a given pixel.

To further enhance the realism of our simulations, we introduce a combination of Poisson noise with a signal-to-noise ratio (SNR) of 5 and Gaussian noise with an SNR of 40 into the generated data. This noise model is representative of the types of noise typically encountered in hyperspectral imaging applications, ensuring that our evaluation accounts for the practical challenges of noise interference.

To evaluate the performance of our proposed method, we employ the Root Mean Square Error (RMSE) criterion, which measures the dissimilarity between the true abundance matrix and the estimated abundance matrix. The RMSE is defined as:

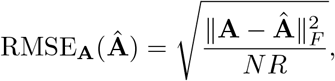

where **A** represents the simulated abundance matrix, which serves as the ground truth, and **Â** denotes the estimated abundance matrix.

With our simulated image data, we found that single-view spectral unmixing (one excitation without photobleaching), exhibited the highest average RMSE Figure 2(e). The multi-view method that incorporates both emission and photobleaching dynamics, showed a reduced average RMSE. For both experimental setups, the RMSEs decreased as the number of excitations increased; however, the multi-view with photobleaching dynamics approach consistently outperformed the approach without inducing and incorporating photobleaching as demonstrated by consistently lower RMSEs.

### 2.2 Real biological images

#### 2.2.1 Imaging strategies

To benchmark BEEP-Learning, we created a reference set of fluorescence in situ hybridization (FISH)-labeled E. coli cells that act as tunable fluorescent standards. With these uniform biological samples, we are able to label cells with virtually any commercially available fluorophore, e.g., any fluorophore that can be covalently conjugated to the 5’ end of an oligonucleotide probe Amann and Fuchs (2008) With these cellular reference standards, we created two kinds of samples, (1) pure populations for extracting BEEP reference signatures and for rigorous quantitative benchmarking of BEEP-Learning against state-of-the-art fewer-view linear unmixing models and (2) mixed communities for qualitative benchmarking of BEEP-Learning on real biological mixed images of unknown composition.

For BEEP data, we employed individual excitations at wavelengths of 445 nm, 488 nm, 514 nm, 561 nm, 594 nm, and 639 nm. This time-modulated, single wavelength excitation strategy is in contrast to simultaneous excitation using fixed combinations of usually 3 laser wavelengths, e.g., 488 nm, 561 nm, and 639 nm, or 445 nm, 514 nm, and 594 nm wavelengths. While any number of discreet excitation wavelengths could be used simultaneously in an imaging setup, in practice, this number lasers can be used simultaneously in epifluorescence microscopy is limited by the recoreded emission wavelength “real estate” blocked by multi-pass dichroic beamsplitters Waters (2009). Importantly, to ensure a relatively fair comparison, both strategies should ideally be applied to the same region; however, due to our intentional imposition of photobleaching, which rendered previously excited molecules incapable of emitting light Song et al (1995), the same region could not be reused. As a result, the two different strategies were applied to different regions, that were qualitatively similar since the samples were homogeneous populations of labeled E. coli cells.

Additionally, we introduced a moderate-multiplexed imaging strategy (MIS), in which two lasers are used simultaneously as a single excitation source. Specific wavelength combinations, 445 nm and 561 nm, 488 nm and 594 nm, or 514 nm and 639 nm, were applied. Pairs of excitation wavelengths were chosen to maximally excite as many fluorophores as possible, e.g., any two lasers used in an experiment are maximally distributed across the visible and near-infrared spectrum. To maintain clarity and focus in the main text, the detailed results of MIS are included in the supplementary material.

#### 2.2.2 BEEP learning for signature extraction

We acquired spectral images of 13 pure populations of *E. coli* samples. Each pure population was labeled with a different version of the EUB338 universal bacterial FISH probe, each version conjugated to a different fluorophore Amann et al (1990). We then learn the emission and bleaching profiles for each fluorophore within its cellular environment.

The spectral signatures for highly correlated fluorophores, e.g., ATTO 620 and Alexa Fluor 633, derived from the single-view method exhibited significant overlap, along with those obtained from multi-view learning with photobleaching only Figure 3 and Supplementary Figures 9-10. In contrast, the signatures obtained using multi-excitation demonstrated significant differences across different wavelength excitations. Notably, for this highly spectrally correlated pair of fluorophores, the BEEP signatures not only differ at 561 nm and 594 nm excitation, but also at 639 nm, where AF 633 decays faster than ATTO 620. We hypothesize that this additional information provided by BEEP signatures underlies the improvement in unmixing results.

**Fig. 3.**
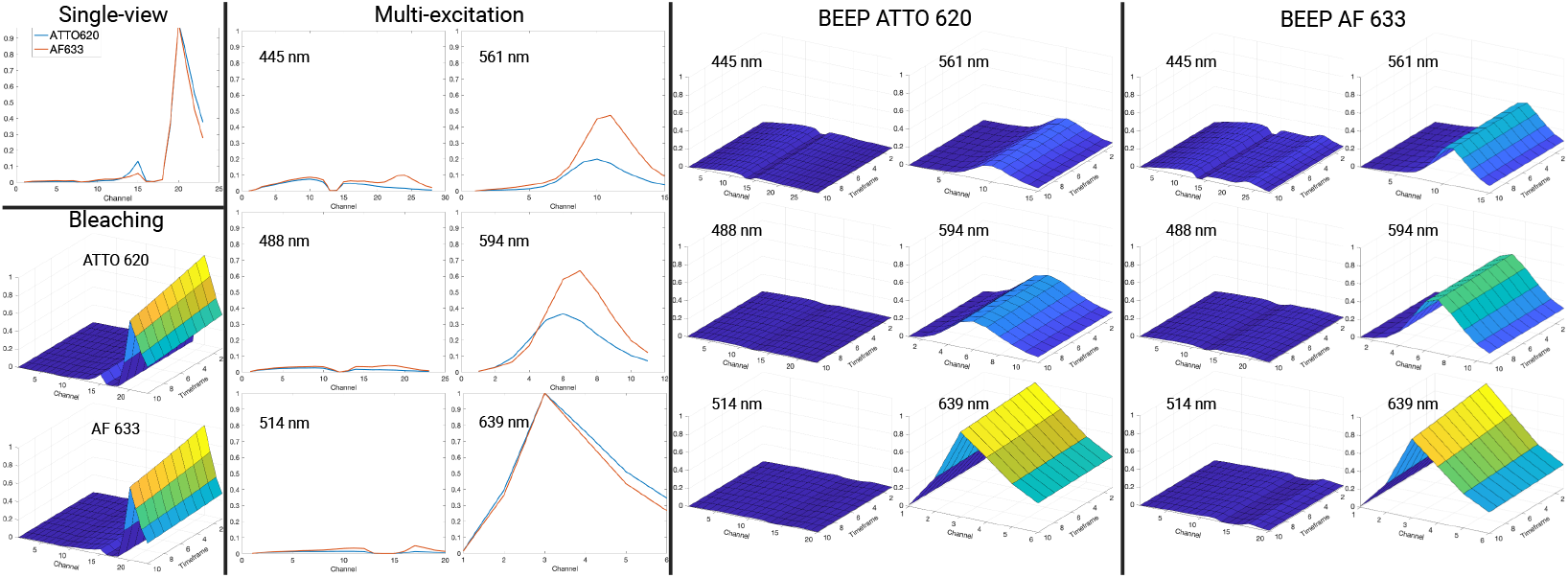
Spectral signatures of ATTO 620 and AF 633 learned using four methods: (1) single-view learning, (2) multi-view learning with photobleaching and emission spectra (bleaching), (3) multiview learning with excitation and emission spectra (multi-excitation), and (4) BEEP.

#### 2.2.3 Unmixing results of reference images

Using these signatures, we estimated the relative abundances of fluorophores. Notably, during the unmixing process, we assumed that all fluorophores were present in each set of images, even though only a single fluorophore was present in each reference image. This assumption enabled us to quantify the accuracy of the unmixing results across different methods, as shown in Figure 4. To qualitatively demonstrate the accuracy in unmixing, we pseudo-colored each pixel in unmixed images according to the fluorophore that was determined to have the largest abundance in that pixel. According to this logic, perfect unmixing results should display a pure color corresponding to the correct fluorophore in all pixels that represent signal from an E. coli cell and the degree to which different colors appear within the same cell approximates the error in unmixing.

**Fig. 4.**
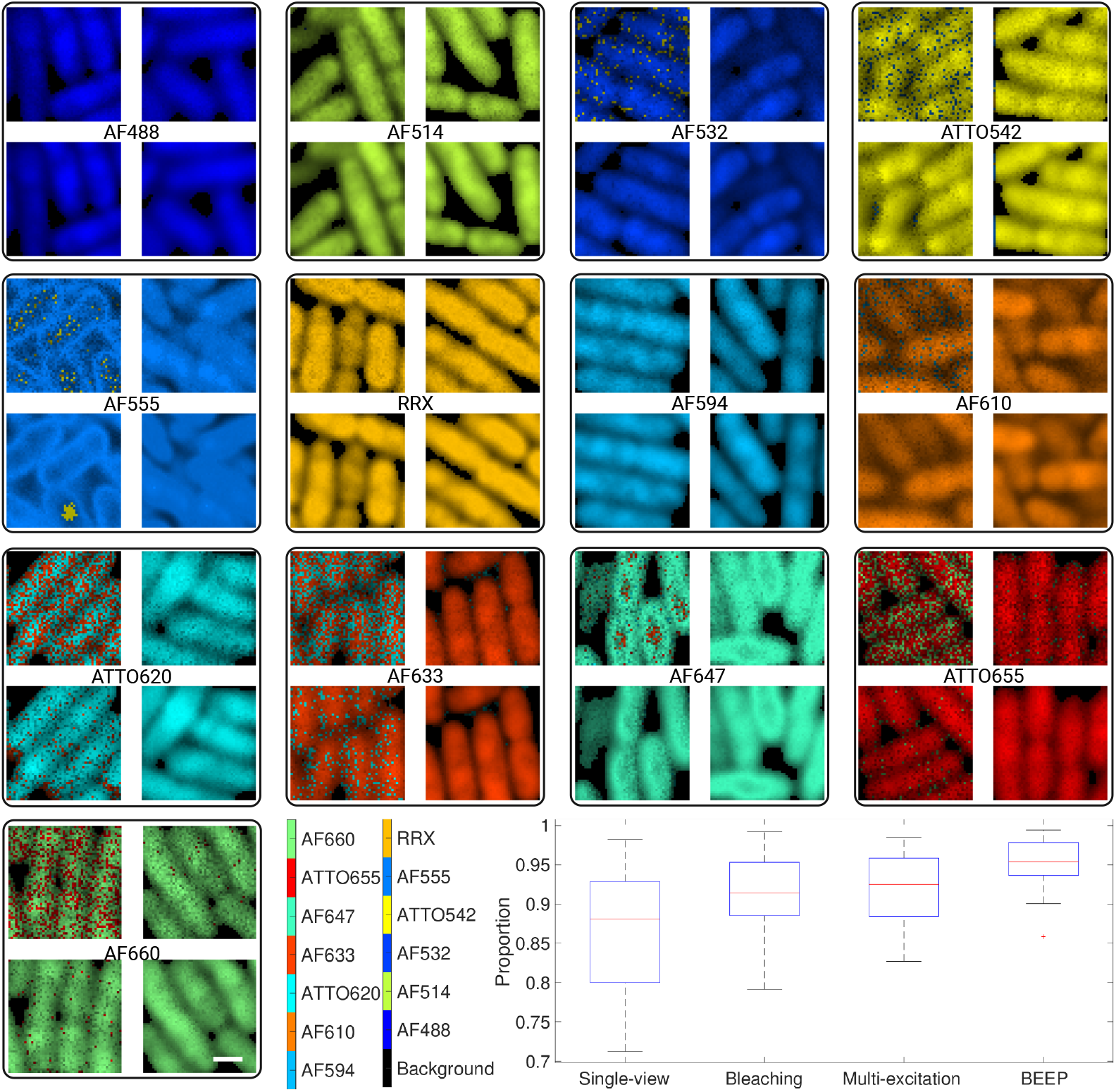
Unmixing results of reference images for AF 488, AF 514, AF 532, ATTO 542, AF 555, RRX, AF 594, AF 610, ATTO 620, AF 633, AF 647, ATTO 655, and AF 660, obtained using four methods: single-view (top-left), bleaching (bottom-left), multi-excitation (top-right), and BEEP (bottom-right). The color of each pixel corresponds to the fluorophore with the largest abundance in that pixel. A white scale bar in the bottom-right corner of AF 660 represents 1 µm. The boxplot in the bottom-right corner compares the average proportion of each fluorophore in its reference image across the four methods defined in (1).

The unmixing results obtained using the single-view method were notably noisy. Those obtained using the bleaching method were less noisy but still displayed signficant errors. In contrast, the results from the multi-excitation method appear homogeneous. In addition to this qualitative image presentation, we further performed rigorous quantitative analysis of accuracy in abundance estimation across the different unmixing methods. To evaluate performance, we calculated the proportion of the estimated abundance of the correct fluorophore in its reference image relative to all 13 abundances, defined as follows. Given an estimated abundance vector **a** = (*a*_1_, *a*_2_, …, *a*_*R*_)^*T*^, where *a*_*r*_, *r* = 1, …, *R*, denotes the estimated abundance of the *r*-th endmember in a pixel from a reference image, the proportion of the involved endmember in this pixel is defined as:

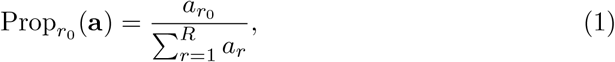

where 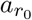 is the abundance of the target endmember. For a reference image, the average proportion of the involved endmember is defined as the mean of these proportions across all pixels. Since each reference image involves only one endmember, the closer the average proportion is to 1, the more accurate the estimated abundance vector. Using a paired *t*-test, we found that the average proportions obtained using BEEP-Learning are significantly higher than those from all other methods Figure 4.

#### 2.2.4 Unmixing results of mixed images

We next benchmarked BEEP-Learning against the three other single or few-view unmixing methods on a real image of a mixture of differently labeled E. coli cells, to simulate a real biological image with complex spatial structure and unknown fluorophore class and abundance Figure 5. The abundance map estimated by the single-view method demonstrated substantial noise, with many cells poorly resolved. By incorporation of photobleaching dynamics into the single-excitation approach, the salt-and-pepper endmember noise was reduced. Similarly, the multi-excitation method improved the endmember resolution of the abundance maps. Notably, BEEP-Learning demonstrated superior performance, achieving the most accurate separation of cells among all methods. These results highlight the benefits of utilizing additional excitation and bleaching-based information to enhance unmixing accuracy.

**Fig. 5.**
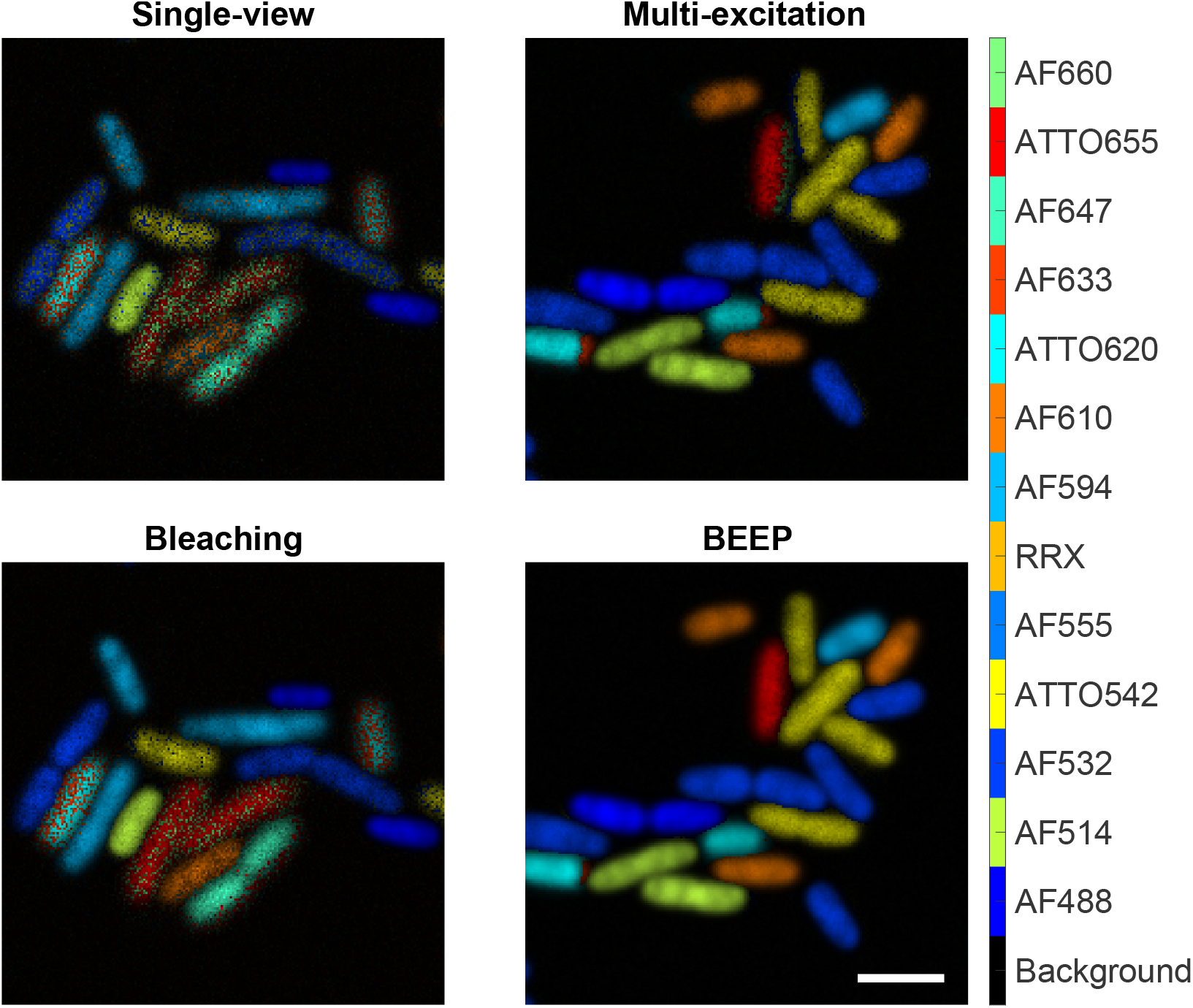
Estimated abundances of a mixed *E. coli* sample obtained using four methods: single-view (top-left), bleaching (bottom-left), multi-excitation (top-right), and BEEP (bottom-right). The color of each pixel corresponds to the fluorophore with the largest abundance in that pixel. A white scale bar in the bottom-right corner of BEEP abundances represents 3 µm.

## 3 Discussion

In this study, we developed and validated BEEP-Learning, a novel framework for spectral unmixing in biological fluorescence microscopy. By integrating information on bleaching, excitation, and emission dynamics, BEEP learning significantly enhances the accuracy of spectral unmixing, addressing a critical challenge in multi-plex fluorescence imaging.

Traditionally, photobleaching has been considered a limitation in fluorescence microscopy, to be avoided as much as possible. This appeals to reason as bleaching causes not only lower signal from fluorophores and therefore lower signal-to-noise ratios, but also because it induces spectral variability when imaging using different excitation wavelengths modulated in the time domain. This nonlinear variability is not accounted for in the linear unmixing least squares model. Our BEEP-Learning framework transforms the photobleaching pitfall into an advantage by leveraging the unique decay profiles of fluorophores as additional information for spectral unmixing. This innovation enables more accurate discrimination between fluorophores with overlapping spectral characteristics, even in complex biological samples.

Through extensive simulations and real-world experiments, we demonstrated that BEEP learning outperforms existing methods, including single-view, bleaching, and multi-excitation approaches. Notably, our method excels in estimating fluorophore abundances in complex biological samples, where high spectral overlap and low signal-to-noise ratios pose significant challenges.

One caveat of BEEP-Learning is that it assumes that the molecular composition of the sample does not change over time. This assumption holds true for fixed samples, e.g., formalin fixed parrafin embedded (FFPE) FISH and immunofluorescence labeled tissues as used in state-of-the-art spatial biology applications Wang et al (2025a); however, this assumption is not valid for live cell imaging with, e.g., fluorescent reporter fusion proteins such as Green Fluorescent Protein (GFP) Miner et al (2024). In addition to employing individual lasers for multiple excitations as a high-multiplexed imaging strategy (HIS), we developed a moderate-multiplexed imaging strategy (MIS), where two lasers are used simultaneously as a single excitation source. This approach achieves a practical balance between the high-multiplexing capability of HIS and the simplicity of conventional imaging methods. Although the unmixing accuracy of MIS is slightly lower than that of HIS, it offers more information than single-view imaging while requiring less experimental and computational time than HIS, making it a suitable alternative for some experimental scenarios, such as live cell imaging where image acquisition time must be minimized to accurately assay systems-level, dynamic molecular interactions.

The flexibility of BEEP learning, which allows for adjustments in parameters such as excitation wavelengths and the number of excitations, makes it adaptable to diverse experimental conditions and imaging hardware setups. This adaptability, combined with its enhanced unmixing accuracy, positions BEEP-Learning as a powerful tool for massively multiplexed fluorescence microscopy.

Here, we evaluated BEEP-Learning in a sample-agnostic manner, meaning only information about the endmembers (fluorophores) used to label cellular structures was used for unmixing. This demonstrates the general applicability of BEEP for different research questions in biological fluorescence. Future work could explore extending this approach in application specific domains; for example, by incorporating cell or sub-cellular morphology or topology information in the unmixing model. Importantly, the tractability of the multi-view machine learning model that underlies BEEP-Learning makes incorporating other “views” of the biological data a straightforward task. This study achieves robust fluorophore discrimination in complex cellular environments and lays the foundation for more sophisticated analyses in biological imaging, where accurate and precise fluorophore discrimination is essential for unraveling complex biological processes.

## 4 Methods

### 4.1 Sample preparation

*E. coli* K12 (ATCC 10798) cells were grown to mid-log phase in Luria-Bertani LB Broth (Difco Laboratories, Inc.). *E. coli* cultures were fixed in 2% paraformaldehyde (EMS Diasum) at room temperature, then stored in 50% ethanol for at least 24 hours before FISH labeling as previously described Valm et al (2011). *E. coli* cells were labeled with the general bacteria probe, EUB338 (GCTGCCTCCCGTAGGAGT) conjugated to a fluorescent dye at the 5’ end.

### 4.2 Imaging

Data collection was performed using Zen Blue Software, which generated CZI files. These files were then converted to TIF files using ImageJ software, and subsequent data processing and analysis were performed in MATLAB. Spectral images were acquired on a Zeiss LSM 980 confocal microscope with a 32-anode spectral detector and a 63x/1.4 NA objective. The images were collected with a spectral resolution of 9.8 nm per channel. Six lasers with different excitation wavelengths 445 nm, 488 nm, 514 nm, 561 nm, 594 nm, and 639 nm. The number of channels for each combination of lasers was determined by the shortest wavelength of the combination, as fluorophores absorb light energy at specific wavelengths and only emit light at longer wavelengths Lakowicz (2006). The exact number of channels for each laser is provided in the Supplementary Table 1.

Our proposed imaging process offers a high degree of flexibility, allowing for adjustments to various parameters as needed. These variables include:

- Excitation wavelength selection
- Number and combination of lasers used
- Number of timelapse frames and time intervals between frames
- Laser power and sequence of excitations

All variables should be consistent across reference and mixed samples.

To minimize the impact of artifacts caused by using different main beam splitters, we employ an intensity-based image registration algorithm Goshtasby (2005) to align the images. This algorithm applies a 2D geometric transformation to optimize the similarity between images of the same field of interest, ensuring accurate registration and minimizing artifacts.

### 4.3 BEEP learning

Each BEEP spectral image sequence consists of five dimensions: excitation dimension, spatial dimensions, emission dimension, and the temporal dimension resulting from photobleaching. The two spatial dimensions were flattened into a single spatial dimension. 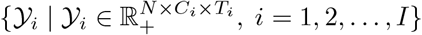 denote the BEEP spectral image sequence recorded at the *i*-th excitation, where *I* represents the number of excitations. The image recorded at the *i*-th excitation has *N* pixels, *C*_*i*_ channels, and *T*_*i*_ timelapse frames. *C*_*i*_ is determined by the laser with the shortest excitation wavelength at the *i*-th excitation.

We assume that the emission profile 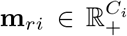 of the *r*-th fluorophore at the *i*-th excitation remains constant throughout the temporal dimension. Additionally, we assume that the bleaching profile 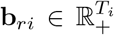 of the *r*-th fluorophore at the *i*-th excitation remains consistent across the emission dimension. Furthermore, we assume that the underlying abundances 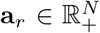 of the *r*-th fluorophore remain unchanged across excitations, temporal dimension, and emission dimension.

The linear mixture model (LMM) has been widely used to analyze multichannel fluorescence images Valm et al (2016); Neher et al (2009); Lakowicz (2006); Chen et al (2022); Gautheron et al (2023); Zimmermann (2005b). LMM assumes that signals from various fluorophores within a pixel combine linearly. To accommodate the unique characteristics of our BEEP spectral image sequence, we introduce BEEP-LMM:

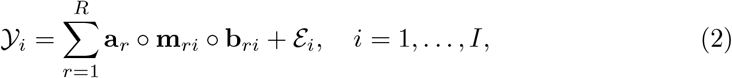

where ○denotes the outer product and ε_*i*_ represents the unknown noise tensor from the *i*-th excitation. In this model, *𝒴*_*i*_ is represented as a rank-one tensor, which is the sum of the outer product of the vectors **a**_*r*_, **m**_*ri*_, and **b**_*ri*_ over all the fluorophores.

To estimate the underlying abundances of all fluorophores in an arbitrary BEEP spectral image, we minimize the noise matrices 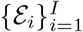 in the BEEP-LMM (2) and solve for both the emission profiles and bleaching profiles simultaneously. This approach is a form of blind source separation Neher et al (2009). However, it faces limitations due to spectral overlap and noise, which can restrict the number of fluorophores that can be accurately resolved. To address this limitation, we adopt a two-stage spectral unmixing approach Rossetti et al (2020). In the first stage, we focus on learning the spectral signatures and bleaching profiles. This is achieved by recording BEEP image sequences of reference samples, each containing a single fluorophore, under different excitation conditions. Using these images in the BEEP-LMM (2), we learn the spectral signatures 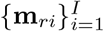 and bleaching profiles 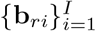 of the *r*-th fluorophore by minimizing the noise matrices. Mathematically, we represent the BEEP reference images of the *r*-th fluorophore recorded at different excitations as 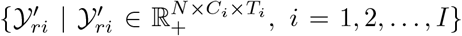 and formulate the optimization problem as follows:

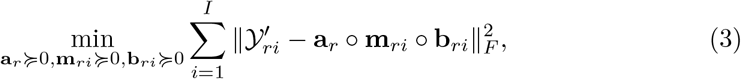

where ∥ · ∥ *F* represents the Frobenius norm of a matrix. By solving this optimization problem (3) for each fluorophore, we can obtain all the spectral signatures and the bleaching profiles, which are then used in the second stage of the unmixing process.

The second stage focuses on estimating the abundances in an arbitrary BEEP spectral image. For the *i*-th excitation, since the spectral signatures and bleaching profiles are already known, we combine them into a single endmember matrix, where each column represents the signature of a specific fluorophore. This can be expressed as follows:

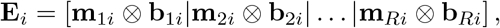

where ⊗ denotes the Kronecker product. To apply 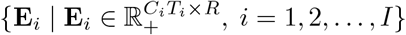 for unmixing, we unfold the 3D images 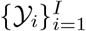 into matrices denoted as 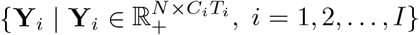 Mathematically, the abundance matrix 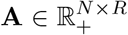 can be obtained by minimizing the noise matrices in BEEP-LMM (2) as follows:

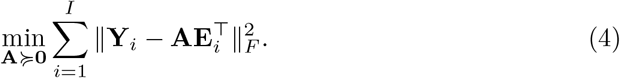

To solve the optimization problem (3), we propose a novel algorithm, outlined in the Supplementary Algorithm, which generalizes the multiplicative update rules Lee and Seung (2000). The optimization problem (4) is equivalent to a nonnegative least squares problem, which can be solved using various algorithms, such as active set method Lawson and Hanson (1995), gradient descent method Johansson et al (2006), and coordinate-wise optimization Franc et al (2005).

## 5 Data Availability

The datasets used in this study are publicly available and can be accessed at the following links:

- https://zenodo.org/records/14210776
- https://zenodo.org/records/14994141

## 6 Code Availability

The source code for this study is available in the GitHub repository: https://github.com/WANGRUOGU/BEEP

## 7 Acknowledgments

This work was supported by National Institutes of Health [grant numbers R01DE031213, R01DE030927, S10OD028600]; and National Science Foundation [grant numbers 1636933, 2111080, 1920920].

## 8 Author information

### 8.1 Contributions

R.W., Y.F., and A.M.V. conceived the method, T.T.H. prepared the samples, R.W. designed the imaging approach, collected and analyzed the data, developed the mathematical model, and wrote the manuscript. R.W., Y.F., and A.M.V. reviewed the manuscript.

