## Supplementary Data File for "BEEP Learning: Multi-View Image Decomposition for Massively Multiplexed Biological Fluorescence Microscopy"

### BEEP Learning for Massively Multiplexed Biological Fluorescence Imaging: Supplementary Material

This supplementary material provides the number of channels for different excitations, the algorithm for BEEP learning, and all signatures and abundance maps obtained through the high-multiplexed imaging strategy (HIS) and the moderate-multiplexed imaging strategy (MIS) described in the main text. For clarity, the multi-excitation and BEEP approaches discussed in the main text are denoted here as multi-excitation (HIS) and BEEP (HIS), respectively. Similarly, MIS without photobleaching is referred to as multi-excitation (MIS), while MIS with photobleaching is denoted as BEEP (MIS).

### 1 Imaging

**Supplementary Table 1** Number of channels with different excitation wavelengths.

| Laser (nm) | 445 | 488 | 514 | 561 | 594 | 639 |
|---|---|---|---|---|---|---|
| Number of channels | 28 | 23 | 20 | 14 | 11 | 6 |

### 2 Algorithm

**Algorithm 1** Multiplicative update rules

**Require:** reference images of the $r$-th fluorophore $\{\mathcal{Y}'_{ri}\}_{i=1}^{I}$
**Ensure:** $\{\mathbf{m}_{ri}\}_{i=1}^{I}$ and $\{\mathbf{b}_{ri}\}_{i=1}^{I}$

unfold $\mathcal{Y}'_{ri}$ into $\mathbf{Y}_{ri}^{(1)} \in \mathbb{R}_{+}^{N \times C_i T_i}, \quad i = 1, \ldots, I$
$\mathbf{Y}_{r}^{(1)} \Leftarrow \left[\mathbf{Y}_{r1}^{(1)} | \mathbf{Y}_{r2}^{(1)} | \ldots | \mathbf{Y}_{rI}^{(1)}\right]$
unfold $\mathcal{Y}'_{ri}$ into $\mathbf{Y}_{ri}^{(2)} \in \mathbb{R}_{+}^{C_i \times N T_i}, \quad i = 1, \ldots, I$
unfold $\mathcal{Y}'_{ri}$ into $\mathbf{Y}_{ri}^{(3)} \in \mathbb{R}_{+}^{T_i \times C_i N}, \quad i = 1, \ldots, I$
initialize $\{\mathbf{m}_{ri}\}_{i=1}^{I}$
initialize $\{\mathbf{b}_{ri}\}_{i=1}^{I}$
**repeat**
  $\mathbf{s}^{(1)} \Leftarrow [\mathbf{m}_{r1} \otimes \mathbf{b}_{r1} | \mathbf{m}_{r2} \otimes \mathbf{b}_{r2} | \ldots | \mathbf{m}_{rI} \otimes \mathbf{b}_{rI}]$
  $\mathbf{a}_r \Leftarrow \mathbf{a}_r \odot \left(\mathbf{Y}_{r}^{(1)} \mathbf{s}^{(1)}\right) \oslash \left(\mathbf{a}_r \mathbf{s}^{(1)\top} \mathbf{s}^{(1)}\right)$
  $\mathbf{s}_{i}^{(2)} \Leftarrow \mathbf{a}_r \otimes \mathbf{b}_{ri}$
  $\mathbf{m}_{ri} \Leftarrow \mathbf{m}_{ri} \odot \left(\mathbf{Y}_{ri}^{(2)} \mathbf{s}_{i}^{(2)}\right) \oslash \left(\mathbf{m}_{ri} \mathbf{s}_{i}^{(2)\top} \mathbf{s}_{i}^{(2)}\right)$
  $\mathbf{s}_{i}^{(3)} \Leftarrow \mathbf{m}_{ri} \otimes \mathbf{a}_r$
  $\mathbf{b}_{ri} \Leftarrow \mathbf{b}_{ri} \odot \left(\mathbf{Y}_{ri}^{(3)} \mathbf{s}_{i}^{(3)}\right) \oslash \left(\mathbf{b}_{ri} \mathbf{s}_{i}^{(3)\top} \mathbf{s}_{i}^{(3)}\right)$
**until** $\sum_{i=1}^{I} \|\mathcal{Y}'_{ri} - \mathbf{a}_r \circ \mathbf{m}_{ri} \circ \mathbf{b}_{ri}\|_F^2$ converges

Note: $\odot$ and $\oslash$ denote element-wise product and division.

### 3 Signatures and Abundances

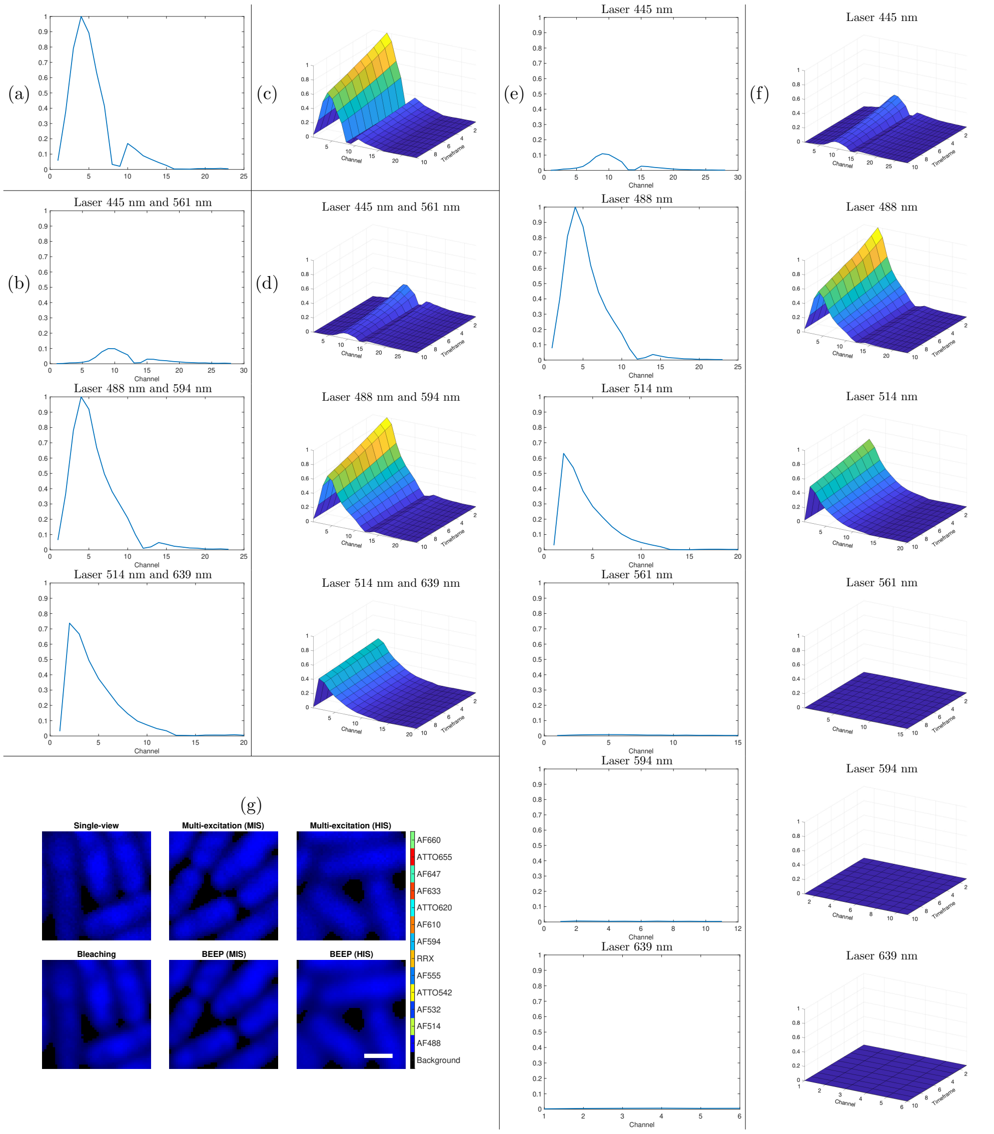


**Supplementary Figure 1** Spectral signatures of AF 488 under (a) single-view, (b) multi-excitation (MIS), (c) bleaching, (d) BEEP (MIS), (e) multi-excitation (MIS), (f) BEEP (HIS), and (g) abundance maps estimated using these methods. In (g), the color of each pixel corresponds to the fluorophore with the highest abundance in that pixel. A white scale bar in the BEEP (HIS) abundance map represents 1 µm.

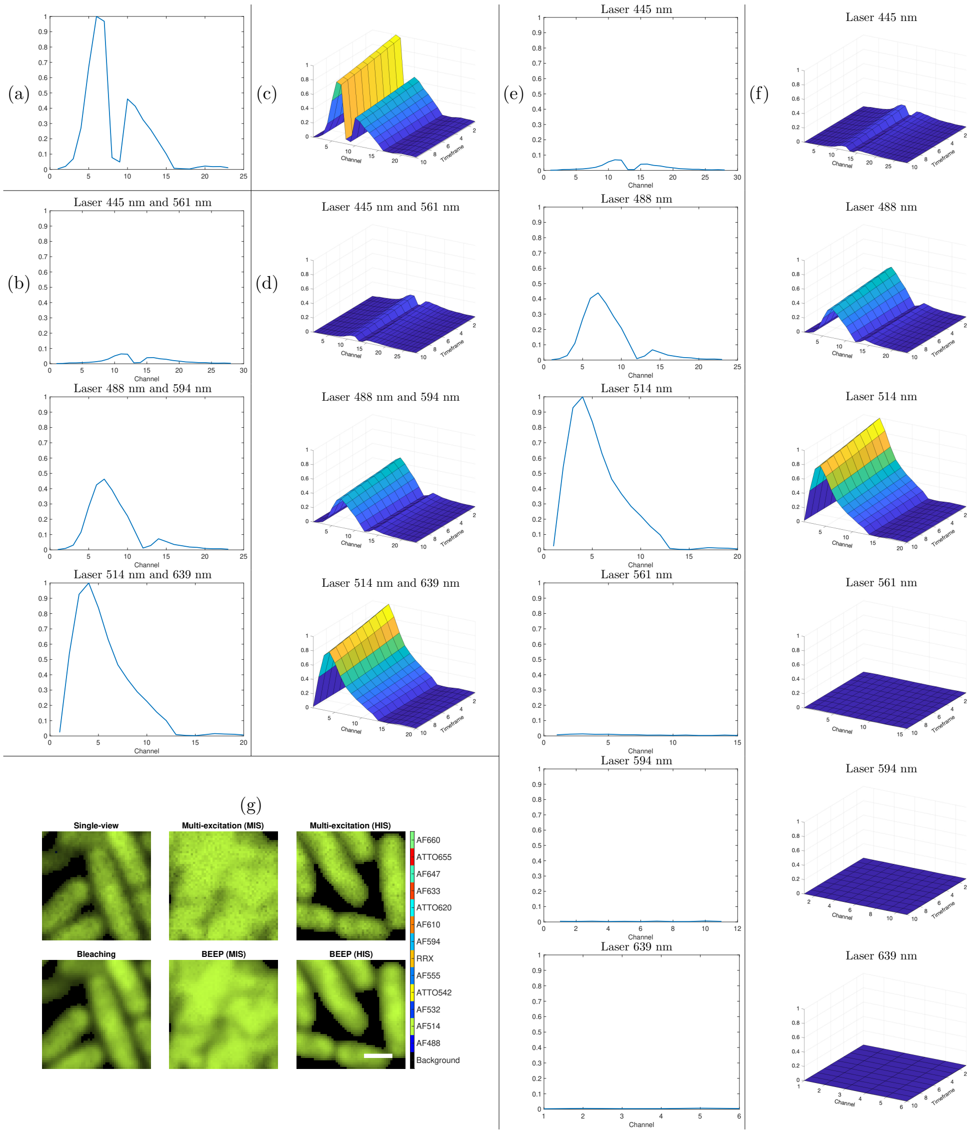


**Supplementary Figure 2** Spectral signatures of AF 514 under (a) single-view, (b) multi-excitation (MIS), (c) bleaching, (d) BEEP (MIS), (e) multi-excitation (MIS), (f) BEEP (HIS), and (g) abundance maps estimated using these methods. In (g), the color of each pixel corresponds to the fluorophore with the highest abundance in that pixel. A white scale bar in the BEEP (HIS) abundance map represents 1 µm.

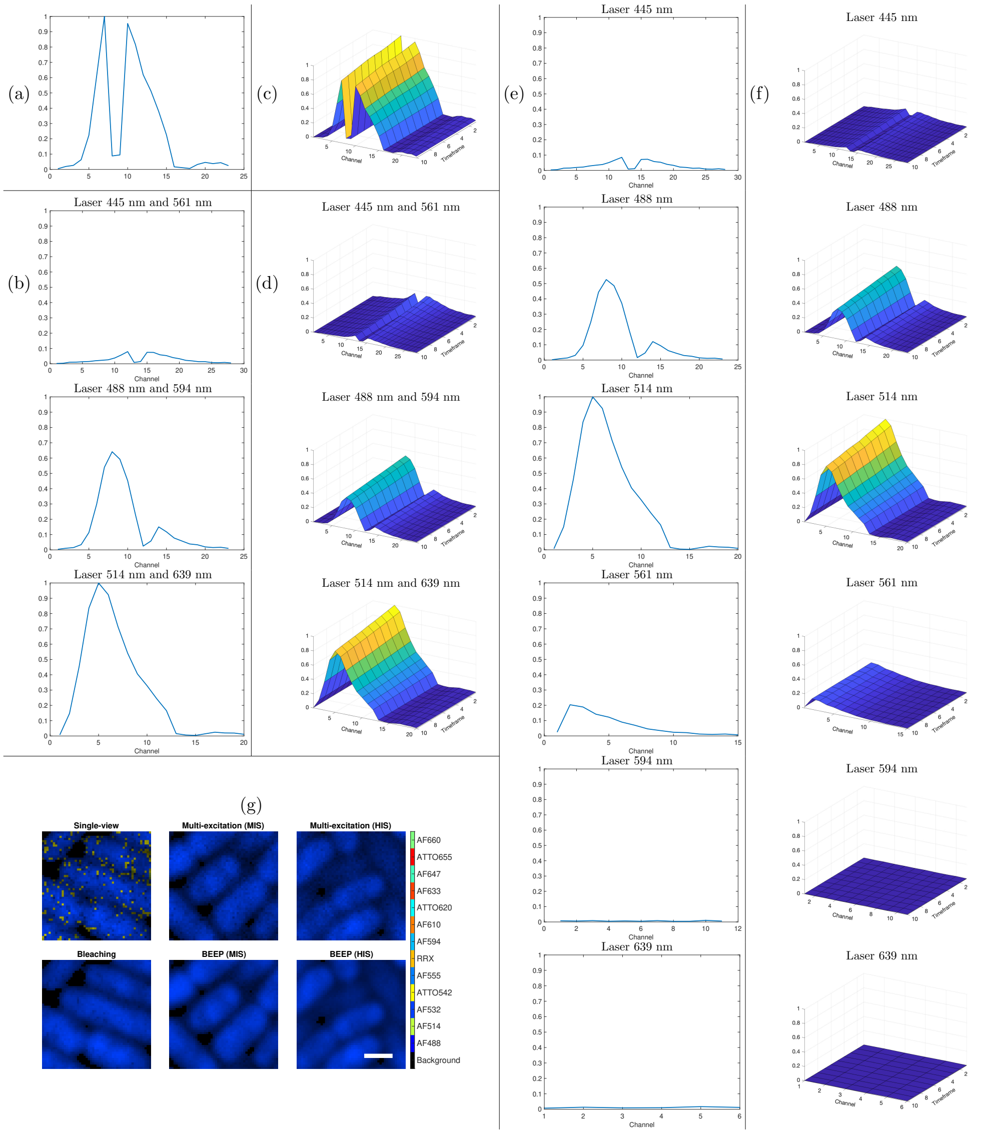


**Supplementary Figure 3** Spectral signatures of AF 532 under (a) single-view, (b) multi-excitation (MIS), (c) bleaching, (d) BEEP (MIS), (e) multi-excitation (MIS), (f) BEEP (HIS), and (g) abundance maps estimated using these methods. In (g), the color of each pixel corresponds to the fluorophore with the highest abundance in that pixel. A white scale bar in the BEEP (HIS) abundance map represents 1 µm.

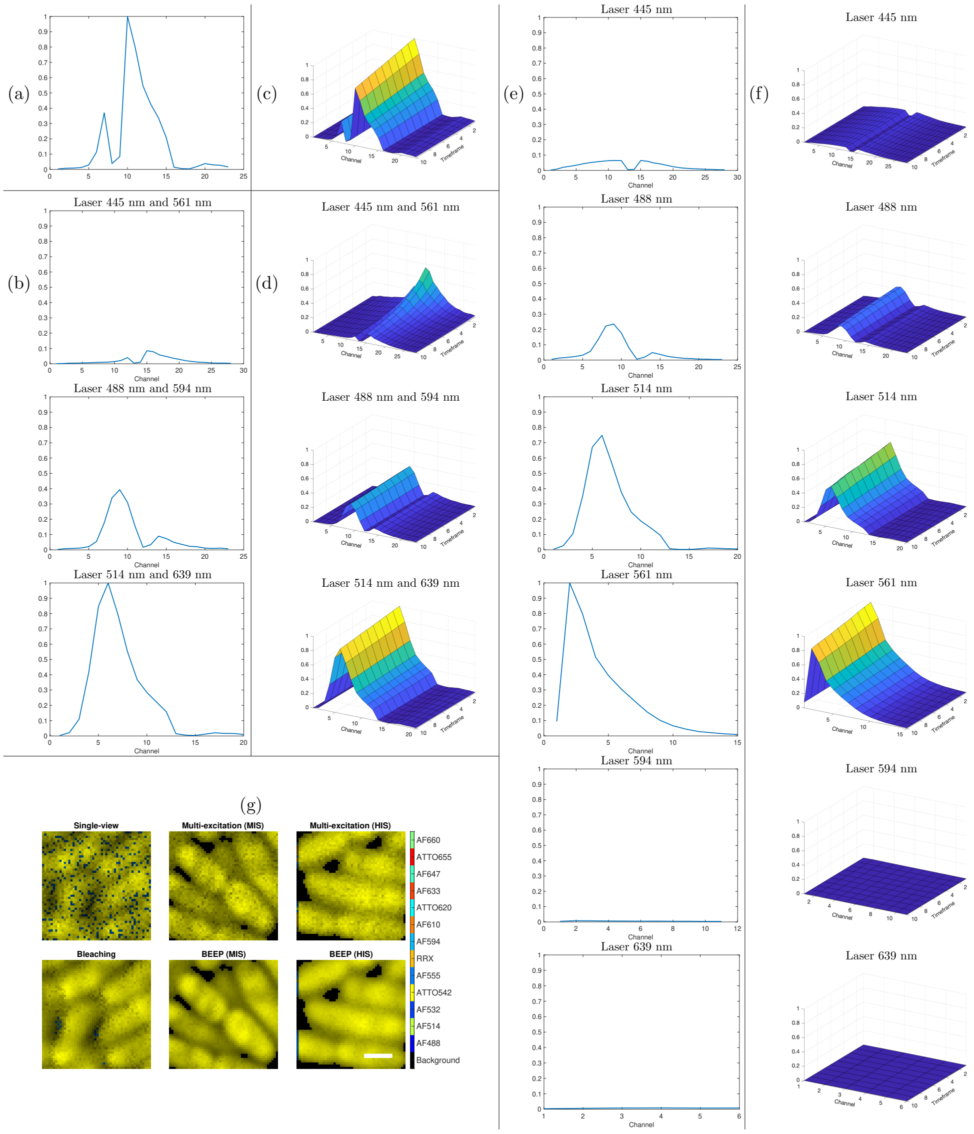


**Supplementary Figure 4** Spectral signatures of ATTO 542 under (a) single-view, (b) multi-excitation (MIS), (c) bleaching, (d) BEEP (MIS), (e) multi-excitation (MIS), (f) BEEP (HIS), and (g) abundance maps estimated using these methods. In (g), the color of each pixel corresponds to the fluorophore with the highest abundance in that pixel. A white scale bar in the BEEP (HIS) abundance map represents 1 µm.

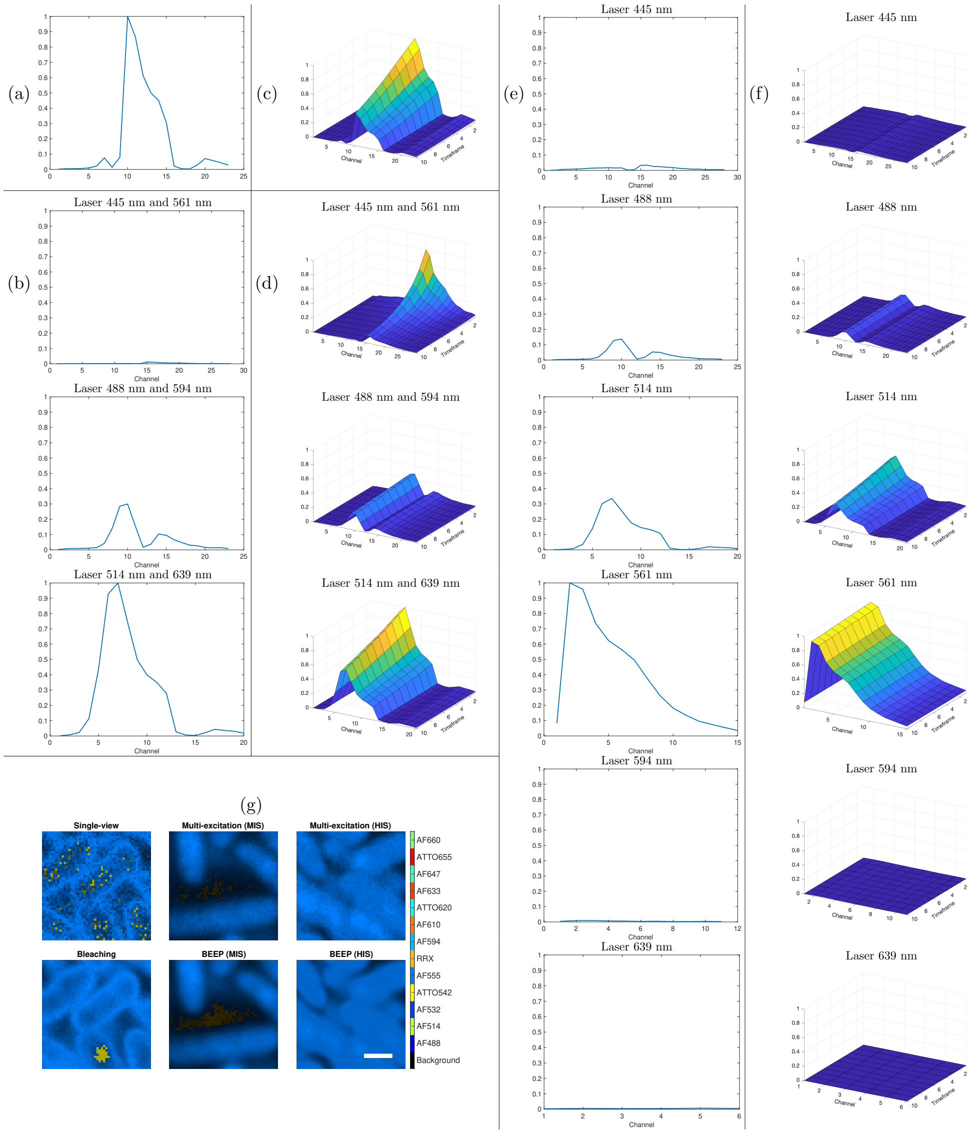


**Supplementary Figure 5** Spectral signatures of AF 555 under (a) single-view, (b) multi-excitation (MIS), (c) bleaching, (d) BEEP (MIS), (e) multi-excitation (MIS), (f) BEEP (HIS), and (g) abundance maps estimated using these methods. In (g), the color of each pixel corresponds to the fluorophore with the highest abundance in that pixel. A white scale bar in the BEEP (HIS) abundance map represents 1 µm.

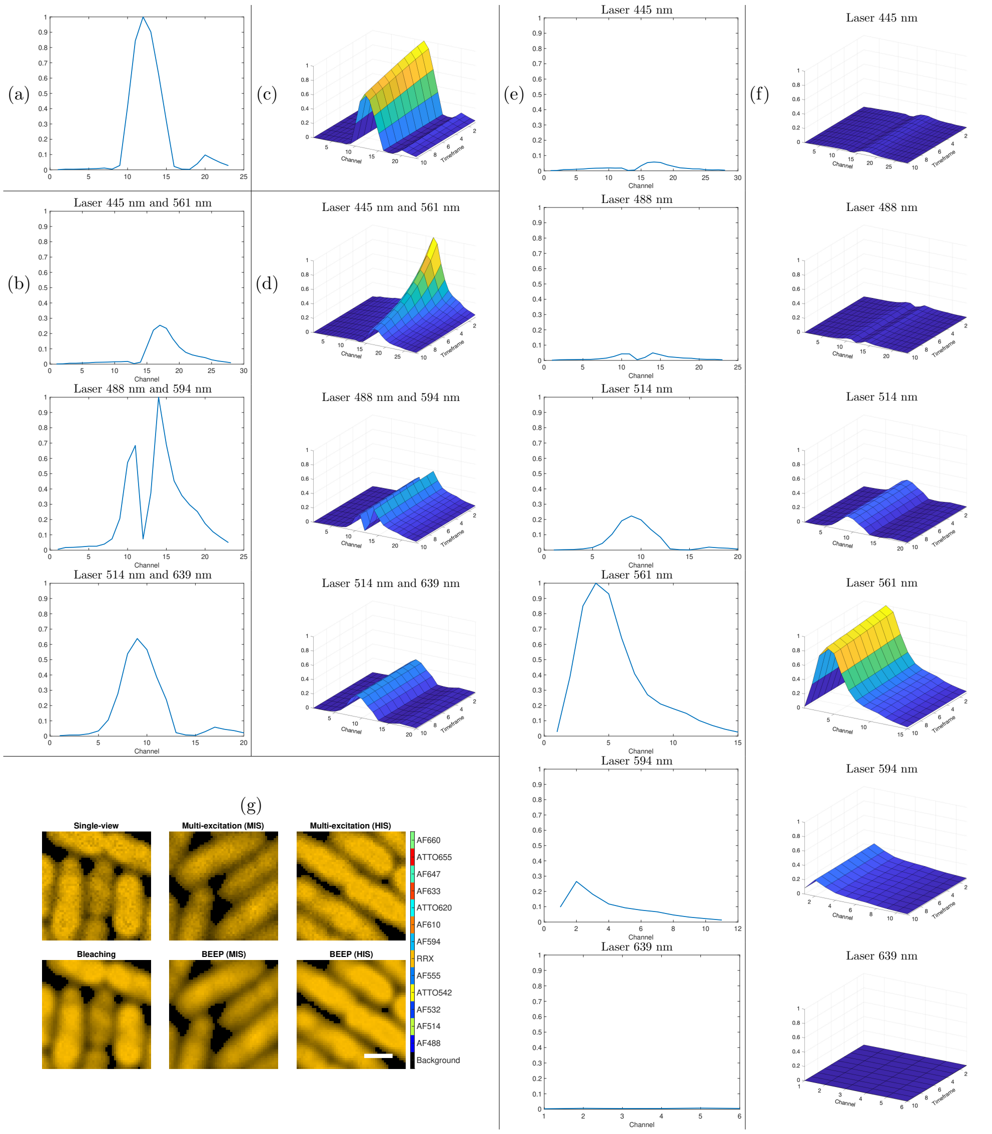


**Supplementary Figure 6** Spectral signatures of RRX under (a) single-view, (b) multi-excitation (MIS), (c) bleaching, (d) BEEP (MIS), (e) multi-excitation (MIS), (f) BEEP (HIS), and (g) abundance maps estimated using these methods. In (g), the color of each pixel corresponds to the fluorophore with the highest abundance in that pixel. A white scale bar in the BEEP (HIS) abundance map represents 1 μm.

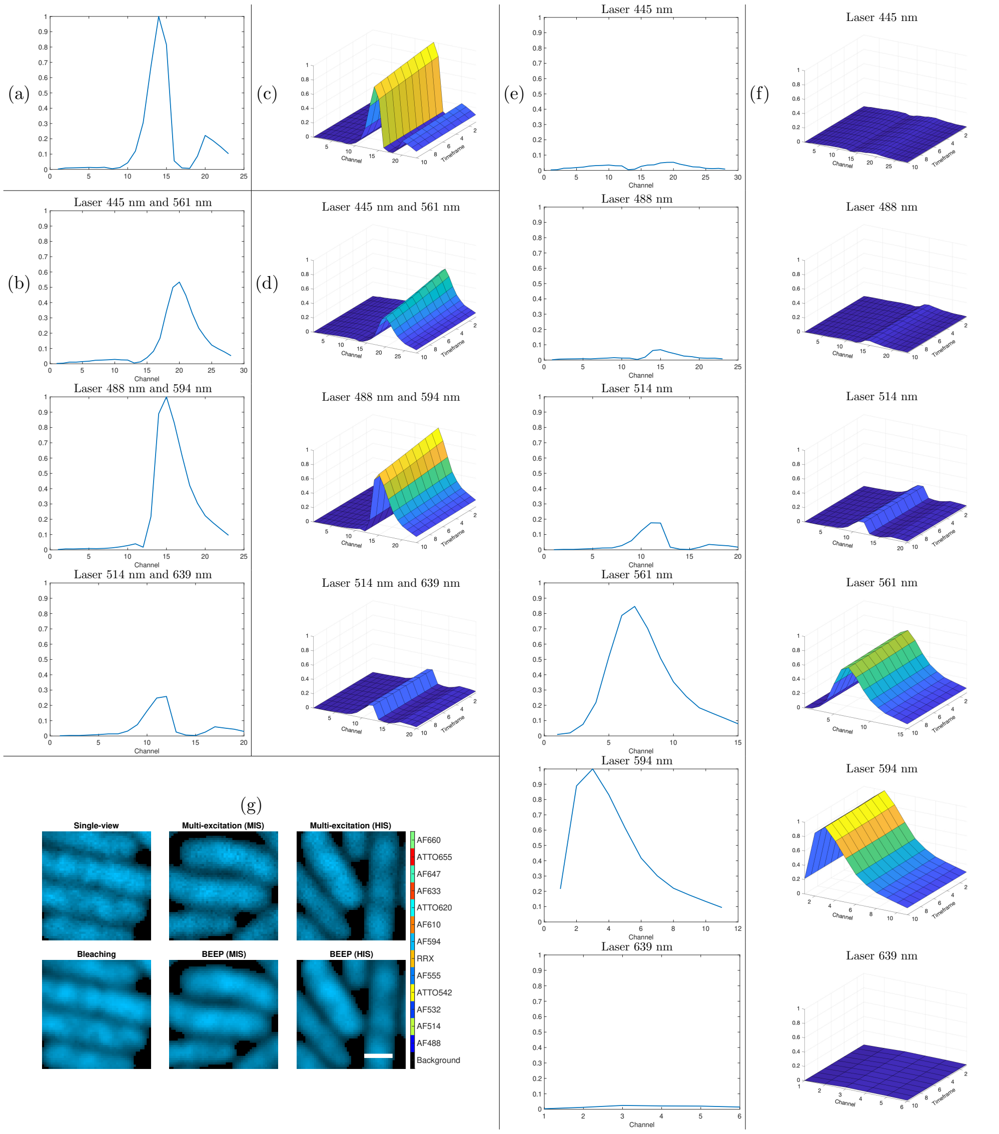


**Supplementary Figure 7** Spectral signatures of AF 594 under (a) single-view, (b) multi-excitation (MIS), (c) bleaching, (d) BEEP (MIS), (e) multi-excitation (MIS), (f) BEEP (HIS), and (g) abundance maps estimated using these methods. In (g), the color of each pixel corresponds to the fluorophore with the highest abundance in that pixel. A white scale bar in the BEEP (HIS) abundance map represents 1 µm.

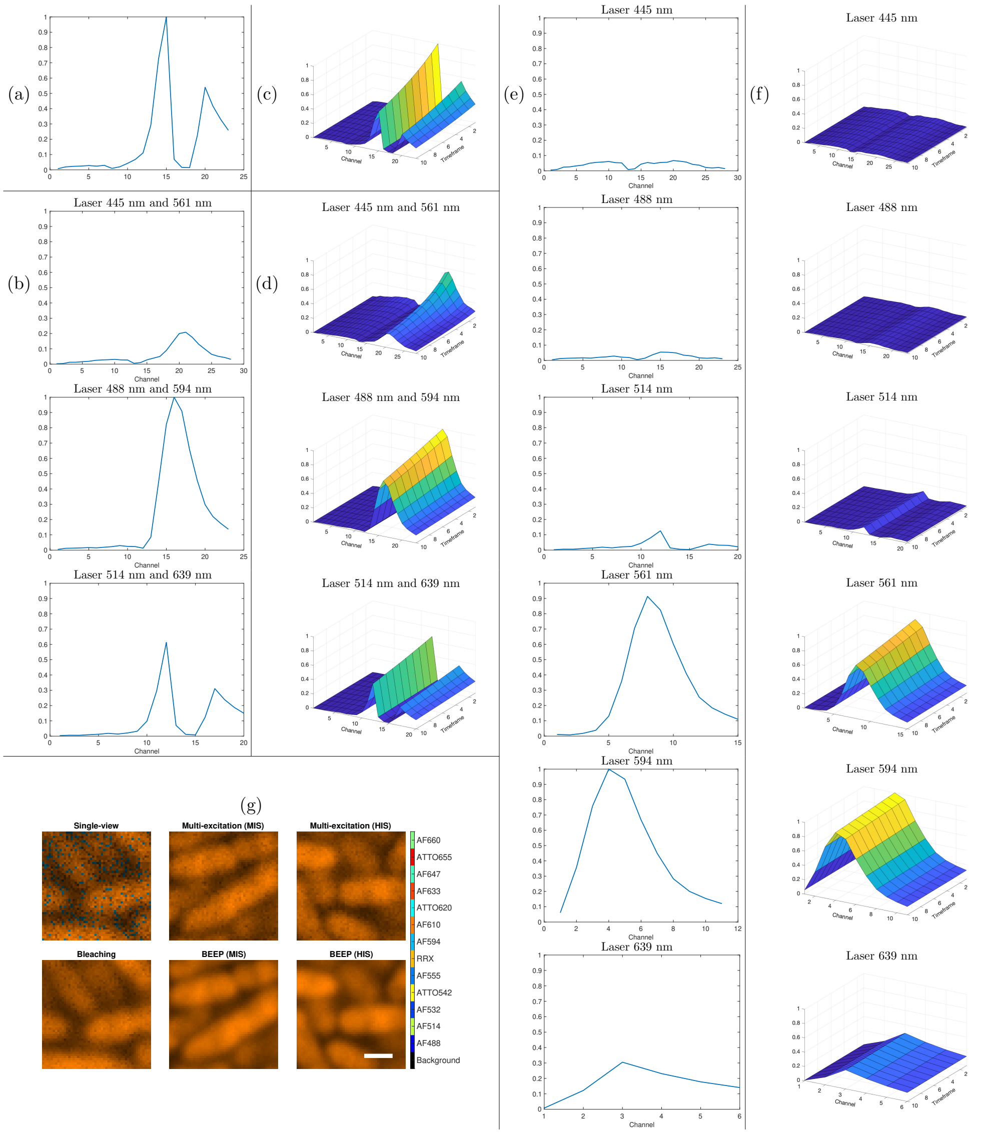


**Supplementary Figure 8** Spectral signatures of AF 610 under (a) single-view, (b) multi-excitation (MIS), (c) bleaching, (d) BEEP (MIS), (e) multi-excitation (MIS), (f) BEEP (HIS), and (g) abundance maps estimated using these methods. In (g), the color of each pixel corresponds to the fluorophore with the highest abundance in that pixel. A white scale bar in the BEEP (HIS) abundance map represents 1 µm.

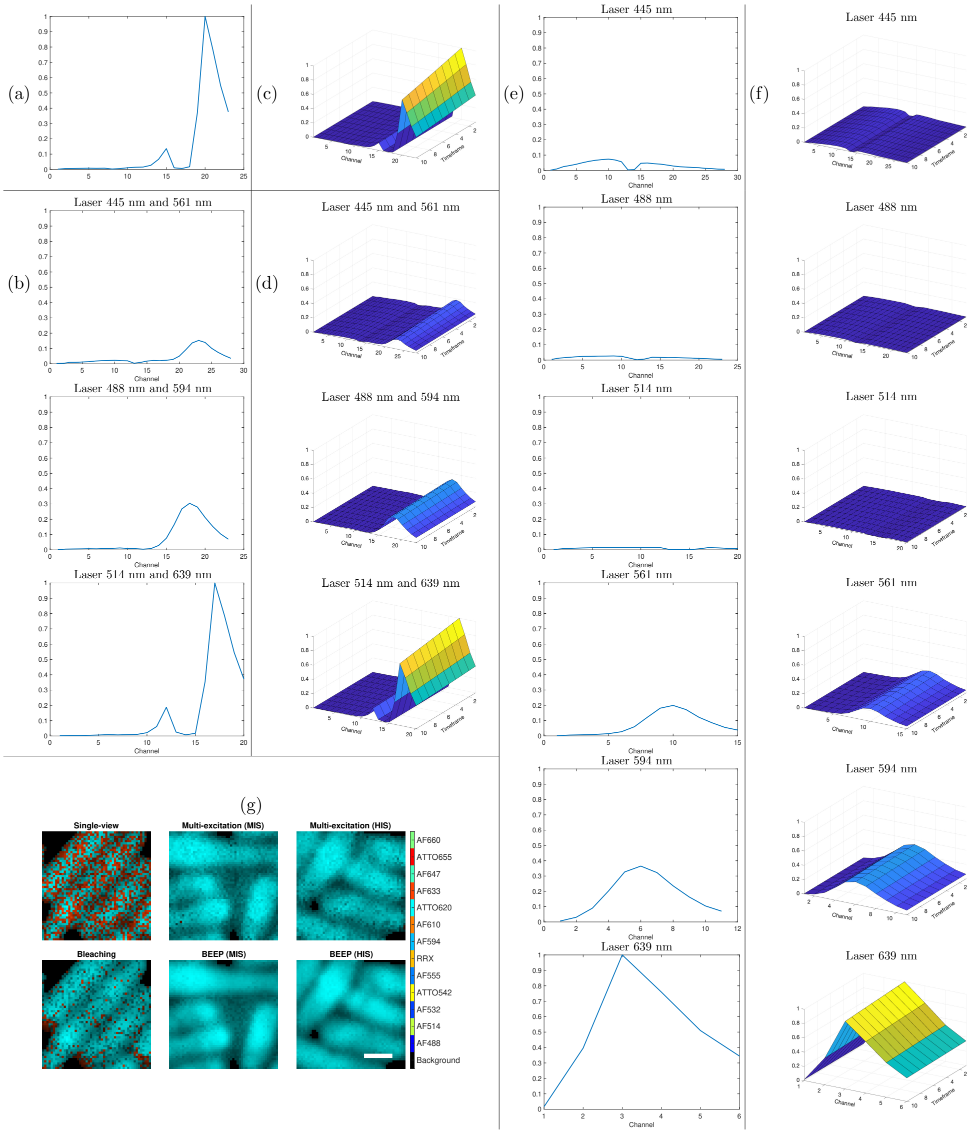


**Supplementary Figure 9** Spectral signatures of ATTO 620 under (a) single-view, (b) multi-excitation (MIS), (c) bleaching, (d) BEEP (MIS), (e) multi-excitation (MIS), (f) BEEP (HIS), and (g) abundance maps estimated using these methods. In (g), the color of each pixel corresponds to the fluorophore with the highest abundance in that pixel. A white scale bar in the BEEP (HIS) abundance map represents 1 µm.

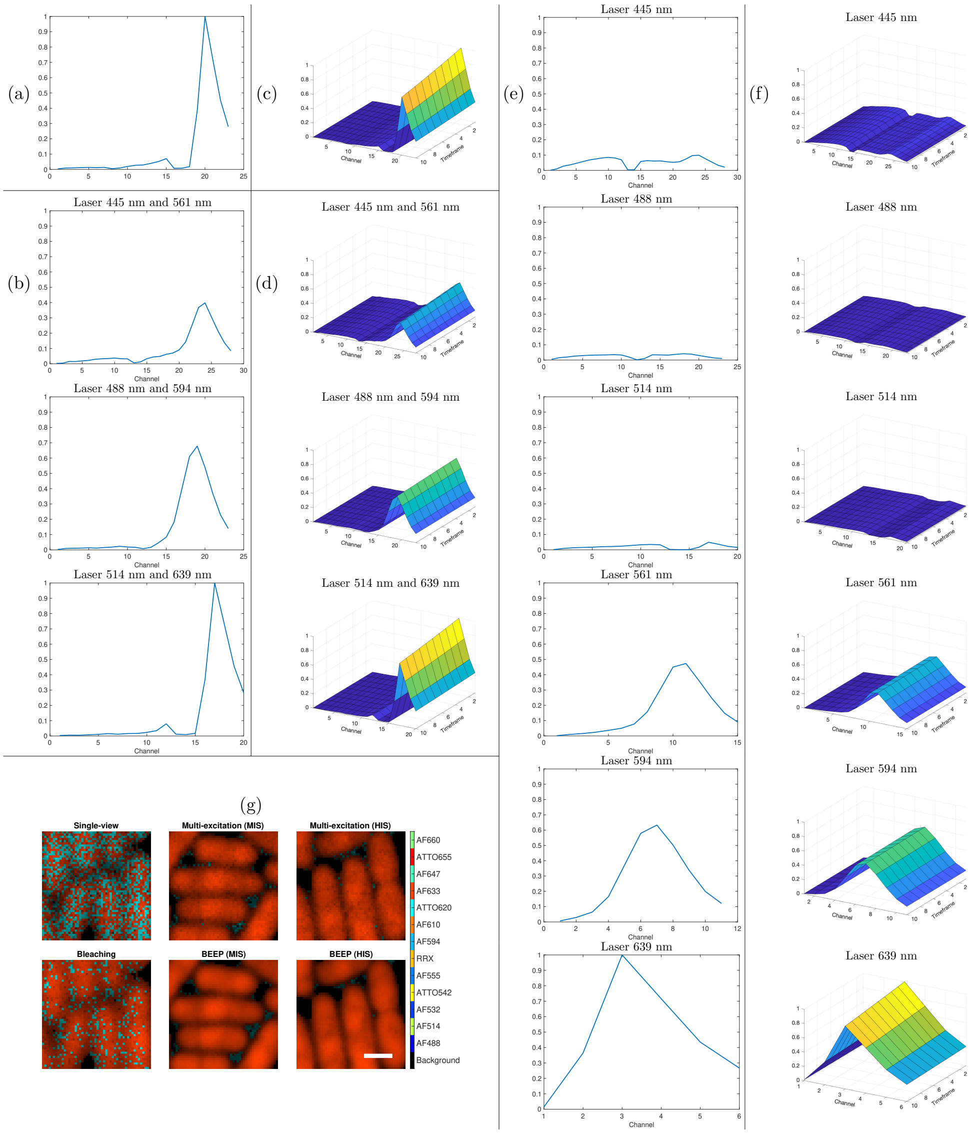


**Supplementary Figure 10** Spectral signatures of AF 633 under (a) single-view, (b) multi-excitation (MIS), (c) bleaching, (d) BEEP (MIS), (e) multi-excitation (MIS), (f) BEEP (HIS), and (g) abundance maps estimated using these methods. In (g), the color of each pixel corresponds to the fluorophore with the highest abundance in that pixel. A white scale bar in the BEEP (HIS) abundance map represents 1 µm.

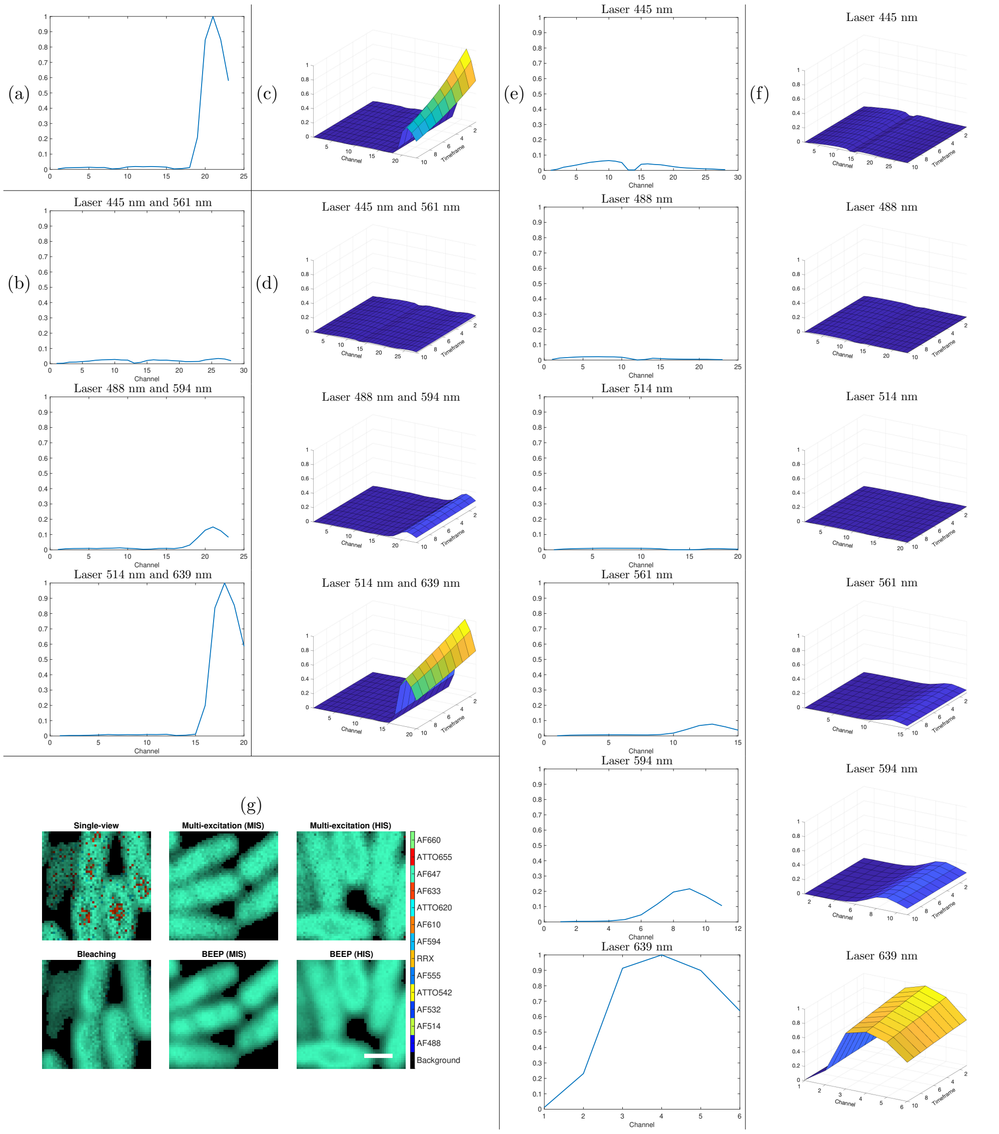


**Supplementary Figure 11** Spectral signatures of AF 647 under (a) single-view, (b) multi-excitation (MIS), (c) bleaching, (d) BEEP (MIS), (e) multi-excitation (MIS), (f) BEEP (HIS), and (g) abundance maps estimated using these methods. In (g), the color of each pixel corresponds to the fluorophore with the highest abundance in that pixel. A white scale bar in the BEEP (HIS) abundance map represents 1 µm.

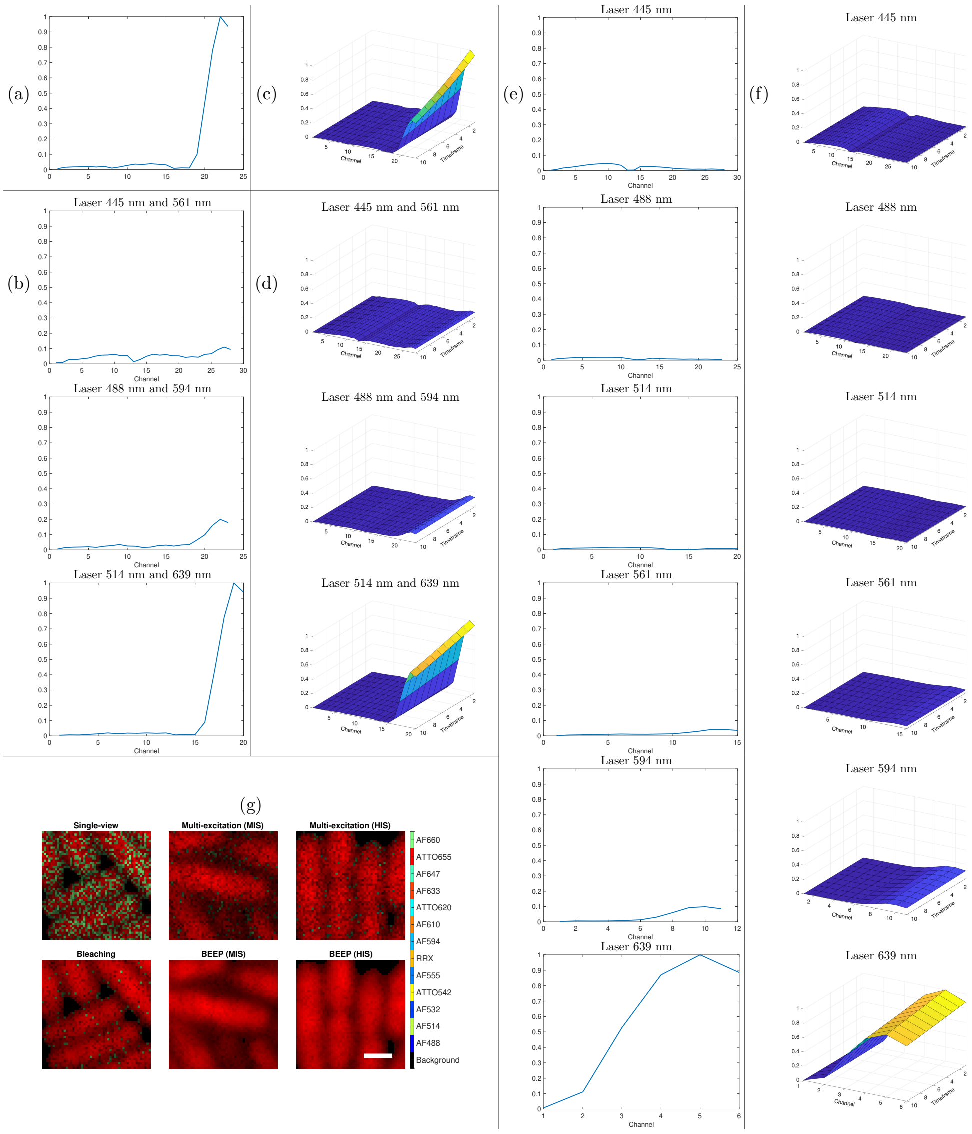


**Supplementary Figure 12** Spectral signatures of ATTO 655 under (a) single-view, (b) multi-excitation (MIS), (c) bleaching, (d) BEEP (MIS), (e) multi-excitation (MIS), (f) BEEP (HIS), and (g) abundance maps estimated using these methods. In (g), the color of each pixel corresponds to the fluorophore with the highest abundance in that pixel. A white scale bar in the BEEP (HIS) abundance map represents 1 µm.

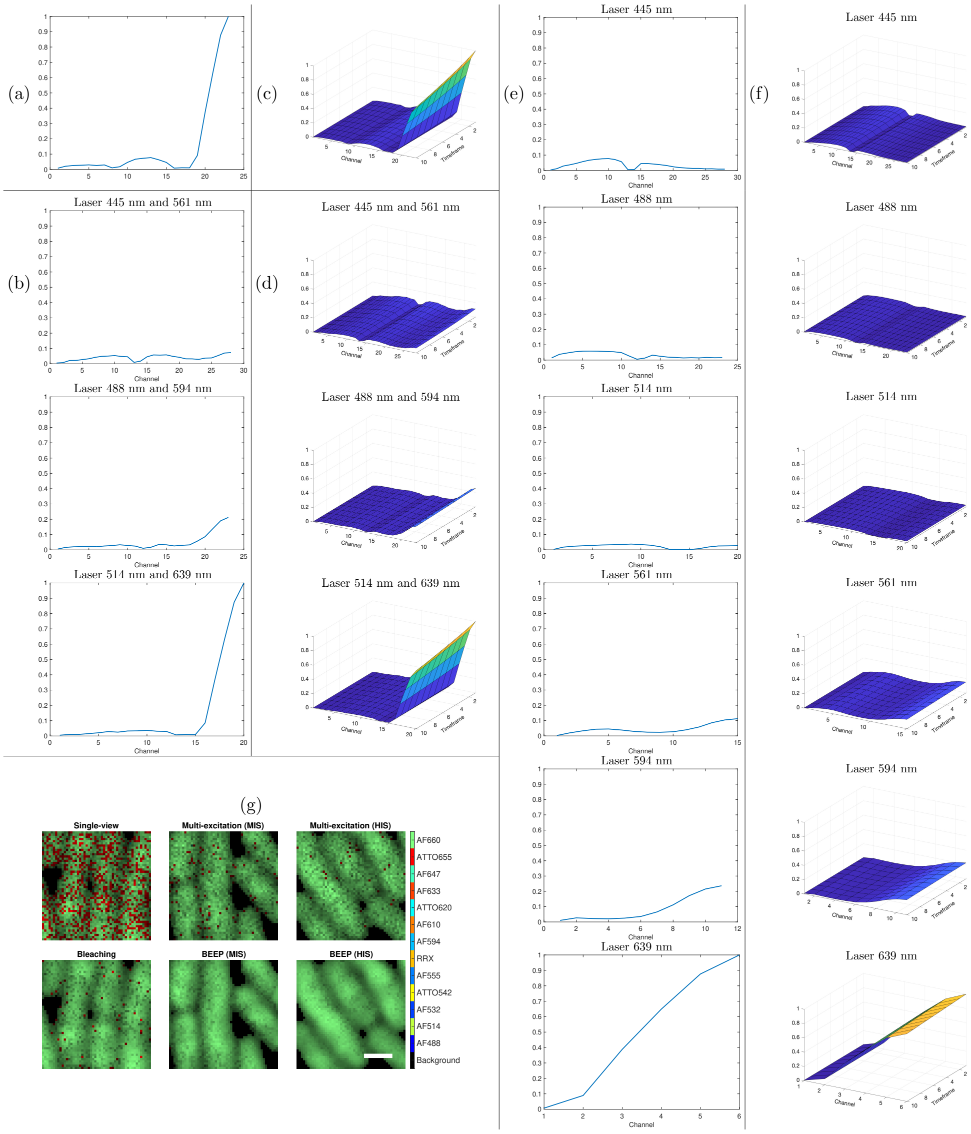


**Supplementary Figure 13** Spectral signatures of AF 660 under (a) single-view, (b) multi-excitation (MIS), (c) bleaching, (d) BEEP (MIS), (e) multi-excitation (MIS), (f) BEEP (HIS), and (g) abundance maps estimated using these methods. In (g), the color of each pixel corresponds to the fluorophore with the highest abundance in that pixel. A white scale bar in the BEEP (HIS) abundance map represents 1 µm.

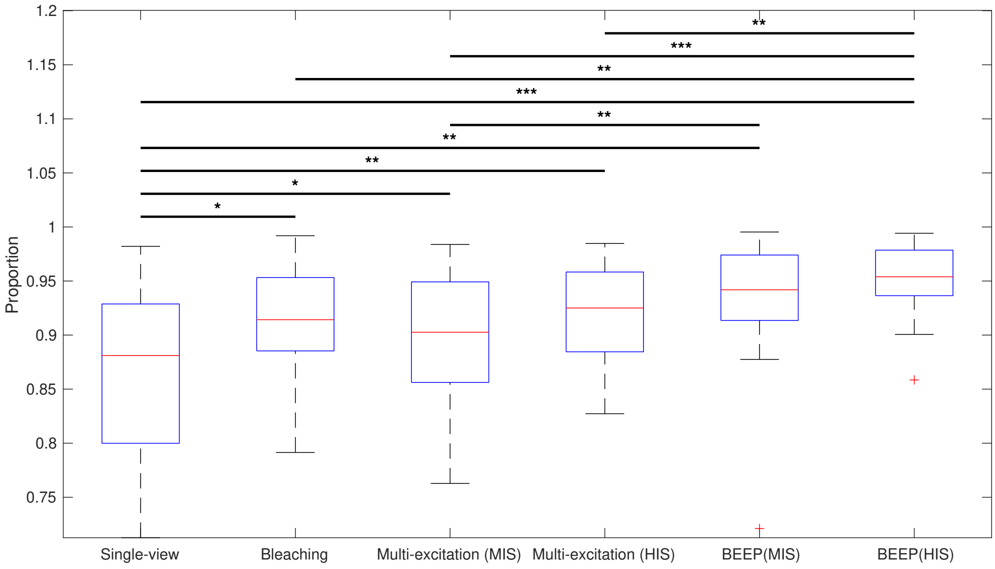


**Supplementary Figure 14** A boxplot comparing the average proportion, defined in the main text, of each fluorophore in its reference image. Statistical significance between pairs of methods was assessed using a *t*-test. The significance levels are indicated by asterisks: * ($p < 0.05$), ** ($p < 0.01$), and *** ($p < 0.001$). Horizontal lines above the boxplots indicate pairs of methods with statistically significant differences.

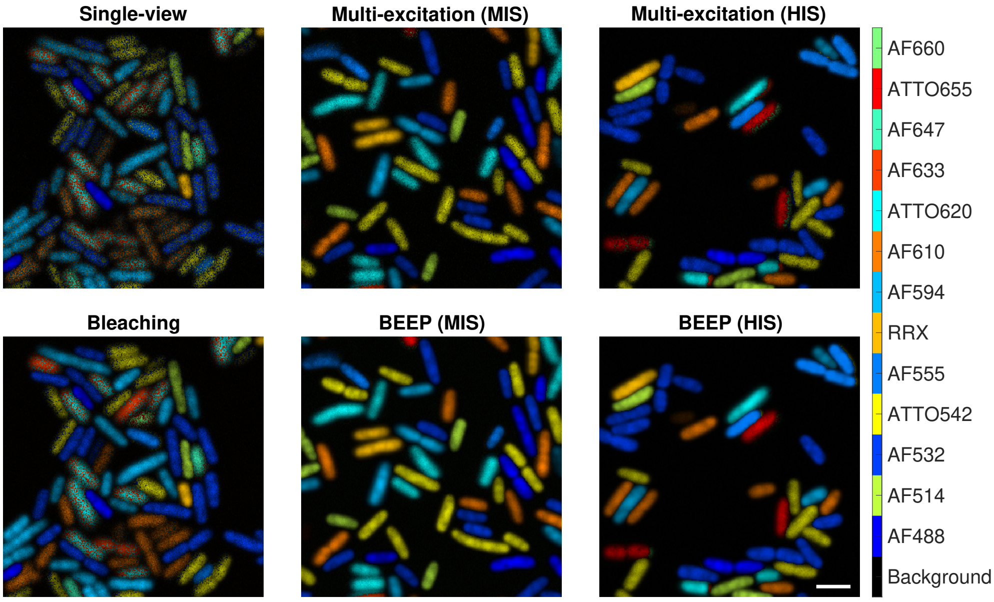


**Supplementary Figure 15** Estimated abundances of a mixed *E. coli* sample. The color of each pixel corresponds to the fluorophore with the largest abundance in that pixel. A white scale bar in the bottom-right corner of BEEP (HIS) abundances represents 3 µm.
